# Differential optogenetic excitation of the auditory midbrain in freely moving behaving mice

**DOI:** 10.1101/2021.02.05.429951

**Authors:** Meike M. Rogalla, Adina Seibert, K Jannis Hildebrandt

## Abstract

In patients with severe sensory impairment due to compromised peripheral function, partial restoration can be achieved by implantation of sensory prostheses for the electrical stimulation of the central nervous system. However, these state of the art approaches suffer from the drawback of limited spectral resolution. Electrical field spread depends on the impedance of the surrounding medium, impeding spatially focused electrical stimulation in neural tissue. To overcome these technical limitations, optogenetic excitation could be applied in such prostheses to achieve enhanced resolution through precise and differential stimulation of nearby neuronal ensembles within the central sensory pathway. Previous experiments have provided a first proof for behavioral detectability of optogenetic excitation in the rodent auditory system. However, little is known about the generation of complex and behaviorally relevant sensory patterns involving differential excitation. In this study, we developed an optogenetic implant to excite two spatially separated points along the tonotopy of the murine central inferior colliculus (ICc). Using a newly-devised reward-based operant Go/No-Go paradigm for the evaluation of optogenetic excitation of the auditory midbrain in freely moving, behaving mice, we demonstrate that differential optogenetic excitation of a sub-cortical sensory pathway is possible and efficient. Here we demonstrate how animals which were previously trained in a frequency discrimination paradigm a) rapidly generalize between sound and optogenetic excitation, b) generally detect optogenetic excitation at two different neuronal ensembles, and c) discriminate between them. Our results demonstrate for the first time that optogenetic excitation at different points of the ICc tonotopy elicits a stable response behavior over time periods of several months. With this study, we provide the first proof of principle for sub-cortical differential stimulation of sensory systems using complex artificial cues in freely moving animals.

## Introduction

Implants restoring auditory function in deaf patients represent the most successful neuronal prostheses. Cochlear implants (CI) alone enabled hearing in over 300,000 patients worldwide^1–3^. Additional electrical approaches stimulate central circuits to restore auditory perception in humans ^4,5^. Electrical stimulation of the central nervous system has, however, limited temporal and, especially, spatial resolution, since electrical field spread impedes spatially focused stimulation^6,7^ and alternative approaches are required to achieve differential stimulation of neuronal ensembles. Optogenetic stimulation (genetically encoded light-activated ion channels) could be applied to neuronal prostheses to achieve more precise excitation^8–11^, resulting in enhanced resolution^6,12,13^. In theory, such differential stimulation should enable the artificial generation of complex percepts.

The behavioral detectability of optogenetic excitation has recently been demonstrated in the rodent auditory pathway^14,15^, the olfactory system^16^, and the primate somatosensory cortex^17^. Differential stimulation has been argued supporting optogenetic stimulation in sensory prostheses, but little is known about the generation of complex and behaviorally relevant sensory patterns. Recent optogenetic studies aimed to ‘mimic’ complex sensory percepts by stimulating superficial cortical neurons. However, the behavioral effects were modest, required complex settings including head fixation and prolonged training periods, and led to increased reaction times^18,19^. Thus, superficial optogenetic excitation of the sensory cortex is inefficient in generating behaviorally relevant sensations. Penetrating, differential stimulation of sub-cortical sensory nuclei may be a more promising approach to generating artificial sensory cues, but requires universally applicable behavioral paradigms for evaluating optogenetic excitation of sensory systems. These paradigms must permit comparing behavior elicited by acoustical and artificial cues, and be stably measureable over several months in the same animal.

Here we demonstrate highly efficient differential optogenetic excitation of a sub-cortical sensory pathway in which the resulting percept reliably drives behavior. Our unilateral optogenetic implant excited two spatially separated points along the tonotopy of the murine central inferior colliculus (ICc). A reward-based operant Go/No-Go paradigm permitted evaluating optogenetic excitation of the auditory pathway compared to sound stimulation in freely moving, behaving mice. We demonstrate for the first time how mice previously trained in a sound frequency discrimination task (1) rapidly generalized between acoustical and artificial cues, (2) generally detected optogenetic stimulation at two separated points of the ICc tonotopy and (3) discriminated between them. Optogenetic excitation at different points of the ICc tonotopy were reliably detected and discriminated over several months. We thus not only uniquely provide a flexible reward-based paradigm for the controlled behavioral evaluation of optogenetic excitation of the rodent auditory pathway, but also the first proof-of-principle for sub-cortical differential stimulation of sensory systems using complex artificial cues in freely moving animals.

## Results

### Freely moving mice detect optogenetic excitation of the auditory midbrain

In the present study, we devised a reward-based operant Go/No-Go paradigm to evaluate optogenetic excitation at different points along the ICc tonotopy of mice and compared the results with those of sound stimulation in the same animals. In short, mice indicated the detection of an auditory target (sound or optogenetic excitation stimuli) by leaving a small pedestal within a response window of 1 s (fig. 1A). To verify the ability of each animal to perform a listening task using complex and continuous discrimination cues, mice were trained in a sound-frequency discrimination task prior to implantation. During surgery, mice were injected with a viral construct into the right ICc (rAAV5-CAG-ChR2-GFP) and implanted with two separated optical outlets, terminating at different depths, 700 µm apart (see fig. 1B-D). After recovery, testing in the sound-frequency discrimination task was repeated to account for habituation and comparable hearing ability post-surgery.

**Figure 1:**
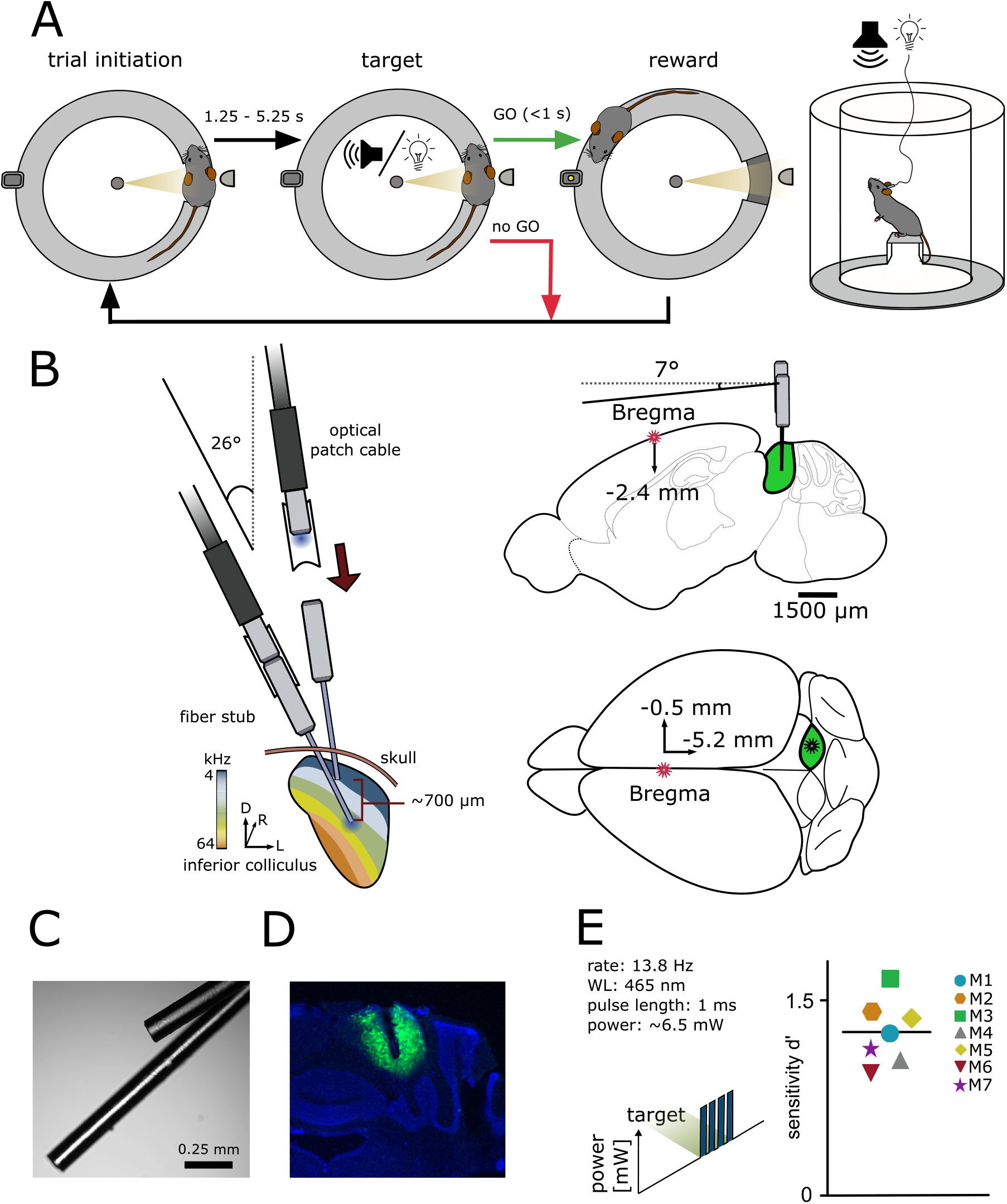
A reward-based operant Go/No-Go paradigm for the evaluation of differential optogenetic excitation of the IC. A: Go/No-Go paradigm. The setup consisted of a circular runway, lined with plexiglas walls and equipped with a platform. The mouse initiated a trial via staying on the platform. After a random delay, the stimulus was presented. If the mouse indicated stimulus recognition by leaving the platform within a time window of one second, a pellet was released as reward at the opposite side of the setup. B: Implantation strategy. Animals underwent a craniotomy followed by injection of the viral construct (rAAV-CAG-ChR2-GFP, 0.9 nl) into the right central inferior colliculus, followed by an implantation of two optical outlets. Optical fibers for LED stimulation with a diameter of 110 µm were used. C: Example image of optical outlets. D: Vector Expression within the IC + DAPI (blue). E: Proof of principle - optogenetic excitation of the IC. Mice performed a paradigm for the detection of optogenetic light pulse trains with a constant power of ∼6.5 mW at either one of the two stimulation points. The overall sensitivity for each animal is shown; black line gives the mean over all animals.

To test for optogenetic excitation of the auditory midbrain, the detectability of a simple cue was measured at one of the two stimulation points in eight animals. The mice indicated the presence of a train of four light pulses (13.8 Hz stimulation rate, 465 nm wavelength, ∼6.5 mW power). Seven test animals reliably detected optogenetic excitation (Chi Squared test), shown as the individual overall sensitivity (fig 5.1E). In contrast to previous studies^18,19^, detection of optogenetic excitation did not require excessive training or habituation, and generalization from sound to light stimulation was mostly rapid (1-2 sessions). One animal failed to generalize (overall sensitivity d’ = −0.1375, three sessions, not displayed). These results demonstrate the suitability of our paradigm to evaluate artificial sensory stimuli.

### Optogenetic excitation can be detected at two well-separated positions within the IC tonotopy

For differential stimulation and establishing complex artificial stimuli in the central circuits, optogenetic excitation at more than one point within the tonotopy is required. We used a psychometric approach to test whether two points can be stimulated separately within the tonotopy of the ICc. Animals had to detect simple stimulus with a varying light level at each stimulation point separately (dorsal and ventral outlet, fig. 2A). All animals completed these tests; optogenetic excitation elicited responses at both stimulation points and the sensitivity increased with increasing light level (fig. 2B). No systematic difference between outlet positions was observed, indicating that excitations at two points within the tonotopy were detectable.

**Figure 2:**
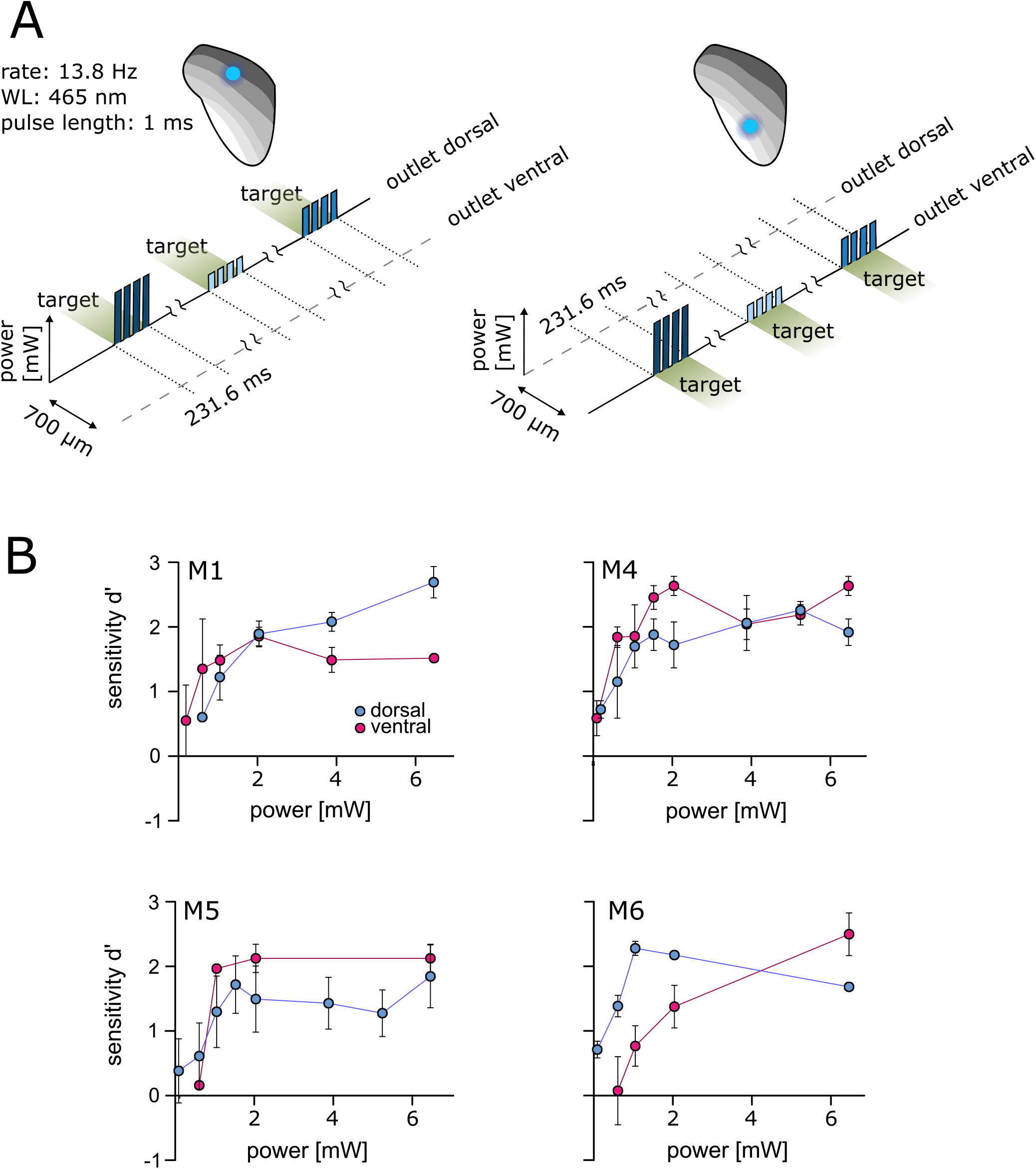
Optogenetic excitation at two well-separated points along the IC tonotopy. A: Psychometric paradigm. To investigate the detectability at two points of excitation, animals performed a simple detection paradigm. The animals had to detect a light pulse train with a varying light level. Sessions were performed for both stimulation points independently. B: Psychometric measurements. The mean d’ at different light levels (absolute power, y-axis, from ∼0.08 to ∼6.5 mW, absolute values) is shown for the dorsal (blue) and ventral (magenta) stimulation point. Error bars are± SEM.

### Mice discriminated between two points of optogenetic excitation along the IC

Next, we investigated whether optogenetic excitation at two points can be used to introduce more complex stimuli by evaluating discrimination between these two points, comparing the results with the post-surgery sound-frequency discrimination task. In the auditory task, the animals indicated a change of frequency within a continuous sequence of tone pips (fig. 3A). Random roving of tone levels between 60 and 66 dB SPL was applied to avoid potential level cues. This task was later mimicked by optogenetic stimulation: animals indicated a change of stimulation point (outlet position) within a continuous sequence of blue light pulses (∼2.2 mW). Stimulation at the dorsal outlet represented the target, whereas stimulation at the ventral outlet served as a baseline (2-point discrimination task, fig. 3B). M1 and M6 immediately performed well in this 2-point task with significant sensitivity (fig. 3C), whereas for M4 and M5 baseline stimulation needed to be adjusted for the mice to successfully discriminate (M5: 50% of background, d’=1.36; M6: 25%, d’=1.98). All background and target light levels were well within mouse detectability (fig. 2B). All animals displayed a similar sensitivity under both conditions.

**Figure 3:**
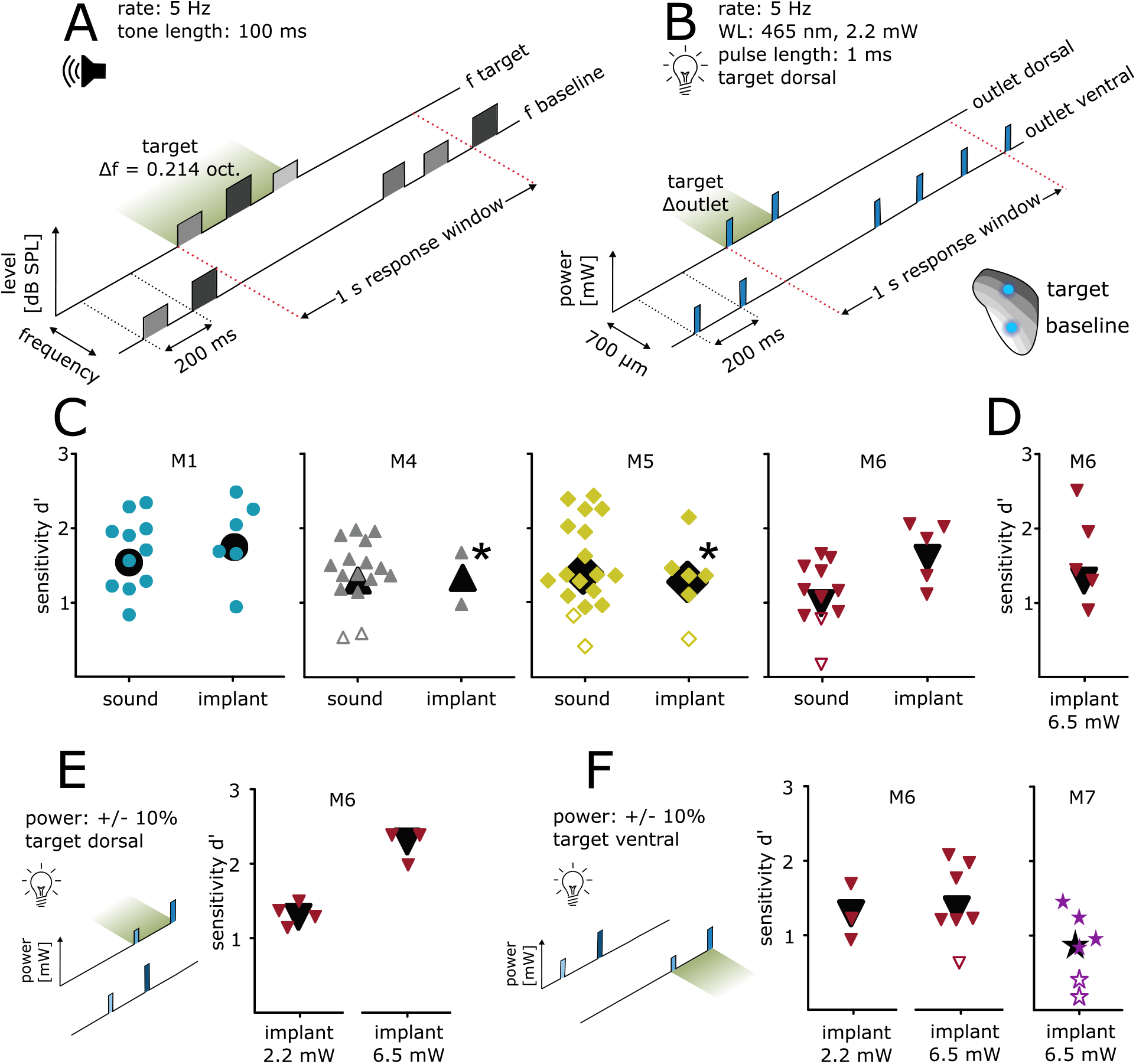
Differential optogenetic excitation of the IC. A: Sound frequency discrimination. Animals had to indicate a change of frequency within a continuous sequence of tone pips (rate: 5 Hz; duration: 100 ms). One session comprised tones of a baseline frequency with a target frequency change of +/- 0.214 octaves. Tone levels were drawn from a uniform distribution between 60 and 66 dB SPL (level roving). B: 2-point discrimination. Animals had to indicate a change in a continuous sequence of light pulses, achieved by the change of the stimulating outlet (465 nm, level: ∼2.2 mW; rate: 5 Hz; pulse duration: 1 ms; distance: 700 µm) C: Behavioral performance. Sensitivity index d’ for all animals and both paradigms is shown. The response towards post-surgery sound (left) and optogenetic excitation (right) is plotted for each individual session, black symbols represent overall sensitivity. If animals did not succeed in discriminating the two stimulation points, the light level of the baseline outlet was reduced (M5: target power: 2.2 mW, baseline power 50% of target level; M4: target power: 2.2 mW, baseline power 25% of target power (upper point) and 50% (lower point). Note that M4 lost the implant afterwards. Open symbols represent sessions with non-significant sensitivity. D: 2-point discrimination at higher light levels. M6 additionally performed behavioral sessions with a power of ∼6.5 mW. E: 2-point discrimination at roving light levels. For M6, the optogenetic paradigms were additionally applied using the same target position but with a roving of power (left: ∼2.2 mW; right: ∼6.5 mW; both ± 10%). F: 2-point discrimination with exchanged stimulation points at roving light levels. These paradigms were repeated but with exchanged target and baseline position (left). Same was applied for animal M7 (right) but at a high power only (∼6.5 mW± 10%).

The light level for the 2-point discrimination task (∼2.2 mW) was chosen as a clearly detectable level (see fig. 2B), but low enough to avoid exciting larger tonotopic areas. To test discriminability dependence on the power, one animal significantly discriminated between the two stimulation points with a level of ∼6.5 mW (M6, fig. 3D), indicating that discriminability did not strongly depend on overall irradiance.

### Discrimination of two excitation points within the IC tonotopy does not rely on level cues and works in both directions

The discriminability thus demonstrated may derive from differences in light delivery (LED/patch cable/fiber stub) or from a difference in opsin expression. To check this, light-level roving was introduced, like the dB-level roving of the sound frequency discrimination task. For two animals (M6 & M7), light pulses of baseline and target were presented at different levels drawn from a uniform distribution of +/-10 %. For one animal, low and high light levels were tested separately (M6, fig. 3E). M6 significantly detected the change in stimulation point immediately in both tasks without additional training or habituation.

Furthermore, the same condition (roving of light level) was applied but with switched target and baseline positions: ventral as target, dorsal as baseline (fig. 3F). M6 discriminated between these for both low (2.2 mW +/- 10 %) and high (6.5 mW +/- 10 %) light levels, but M7 only performed well with the high light level. Although M7 performed poorer than M6, its overall sensitivity was significant (fig. 3F). Thus discriminability is independent of target position and overall irradiance.

### Mice respond faster to optogenetic excitation than to sound stimulation

To better understand the percepts elicited by the different stimuli, we investigated reaction times (RT). The median RT was 73 ms shorter for discrimination of optogenetic excitation than for sound frequency (average RTs: 464 vs. 390 ms, fig. 4A). There was, however, a difference in the temporal stimulus patterns; the sound was 3 tone pips (100 ms/pip), whereas the 2-point excitation target was 2 x 1 ms pulses. We thus evaluated RTs for stimulus detection (see fig. 1E) using a subset of three mice that performed an additional click-detection task using the optogenetic temporal pattern (rate: 13.8 Hz; 4 x 1ms clicks). Again, optogenetic excitation was detected 61 ms faster than sound stimulation (average RTs: 392 vs. 331 ms; fig. 4B).

**Figure 4:**
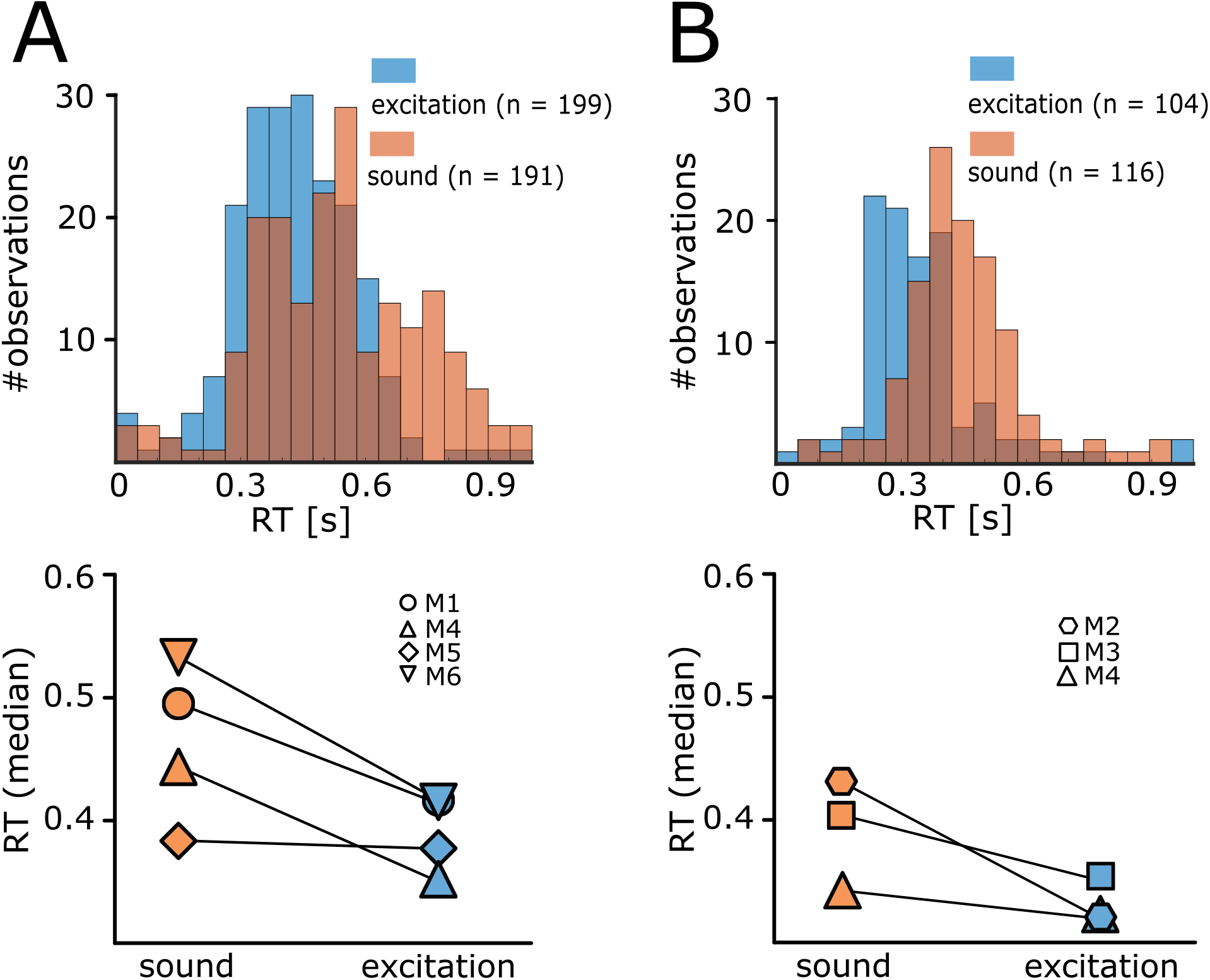
Response Latency’s for sound stimulation and optogenetic excitation. A: Reaction times: Discrimination. Upper panel: example histogram for reaction times (x axis) during optogenetic excitation (blue) vs. sound stimulation (orange) for one animal. Numbers of observations (y axis) were binned with a width of 50 ms. Lower panel: median reaction times (right) in both paradigms (results of C, all animals). B: Reaction times: Detection. The same is shown for the click detection stimulus (orange) and detection of the excitation (blue).

### Optogenetic excitation is not detected visually

To avoid possible visual detection of light stimulation, a ceiling blue-light LED strip continuously pulsed at the stimulus rate during each session. Although this light should mask visually detectable blue light pulses, the possible detection of visual cues from fibers or connectors remains. Thus in all animals that completed the 2-point discrimination, a control task of a set of five independent sessions to detect a light-pulse train in either one of the two outlets with 50/50% probability was conducted (see fig. 1E). In every second session, black foam blocked the implant’s connector (fig. 5A). For open outlets (green), mean hit rates remained relatively stable independent of outlet position (dorsal or ventral). In contrast, the hit rates dropped to chance level for both outlets when blocked (red) but reemerged during the following session (fig. 5B), indicating that the light stimulation was not visually detectable.

### Detection of optogenetic excitation requires the expression of ChR2

Independently of light-sensitive ion channels, thermal effects of light delivery might elicit a neuronal response and the detectability of light stimulation^20^. To control for the necessity of the opsin, 2 animals were trained, transfected, and implanted as previously described, but were injected with a control construct (rAAV-CAG-GFP, fig. 5C; cM1, cM2). Compared with the experimental group, all animals significantly detected a change in sound frequency given as overall sensitivity (fig. 5C & D, left panel). During the light-detection task (compare fig. 1E), experimental animals detected optogenetic excitation, whereas stimulation failed to elicit a significant response in control animals (fig. 5C, D right panel & E). To prove that this was not based on reduced generalization but on the absence of an observable target, light pulses were exchanged with broadband clicks, using the same temporal properties as the optogenetic detection stimulus. Control animals immediately detected click stimuli significantly in the first session and significant overall sensitivity for six consecutive sessions (overall d’, fig. 5C), indicating that the insensitivity of the control animals was not based on a lack of generalization.

**Figure 5:**
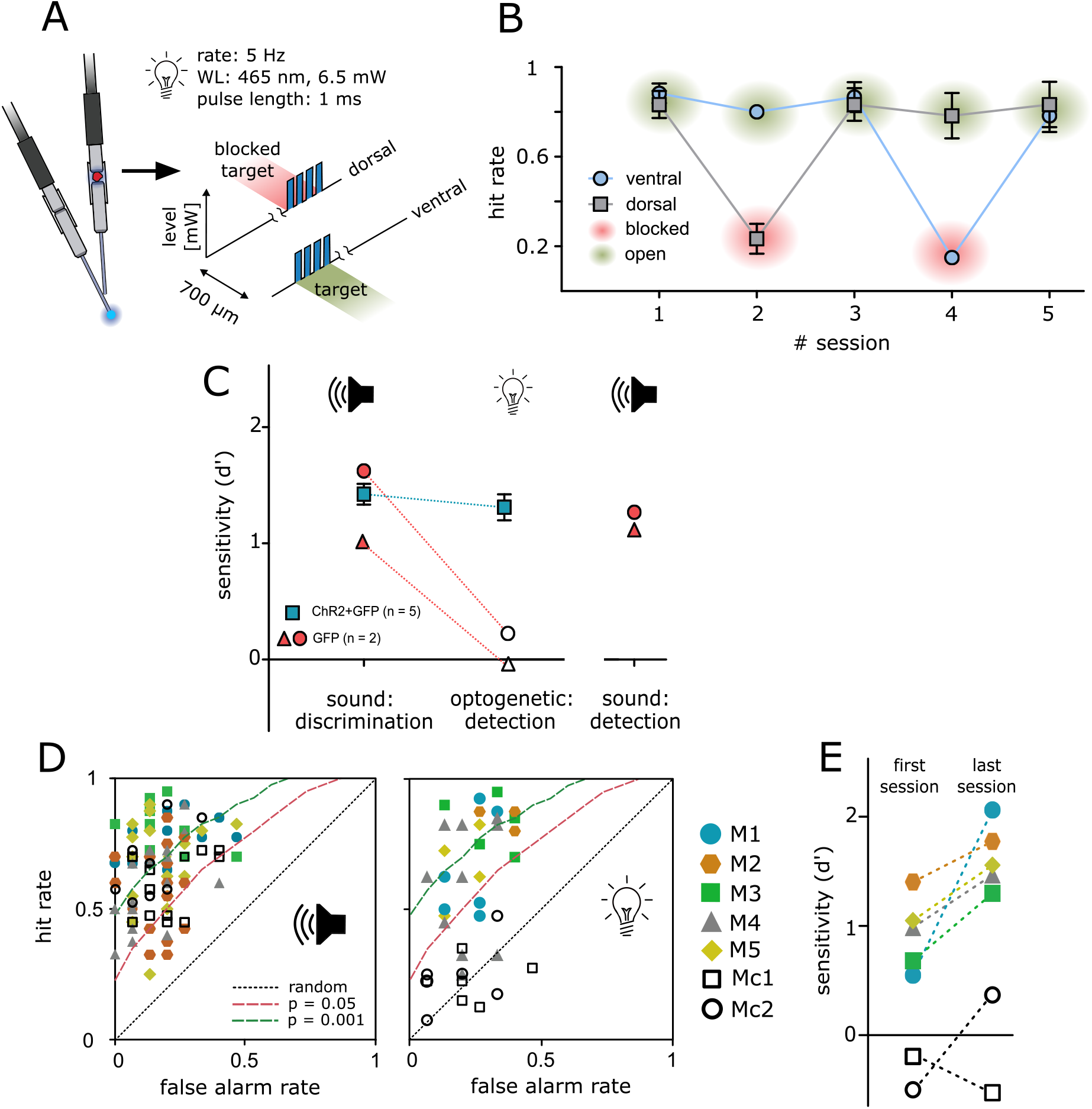
Control experiments. A: Control paradigm. To control for a possible visual light detection, animals performed a detection paradigm with a constant light level (6,5 mW) in which they had to detect a short pulse train as target in either one of the two outlets. In each second session, one outlet was blocked (red) at the connector from patch cable to animals implant. B: Behavioral performance. Mean hit rates (y-axis) are shown for both outlets and five consecutive sessions (x-axis) for animal Ml/5/6. Error bars are ± SEM. C: Control animals. Two animals (grey circles) were injected with a sham virus without the construct for ChR2 (rAAV5-CAG-GFP). Their behavioral performance (sensitivity d’, y axis) is shown in comparison to the experimental group (blue squares, mean ± SEM) for sound frequency discrimination (post-surgery) as well as for the light detection paradigm. To evaluate if the reduced performance of control animals in the detection paradigm for optogenetic excitation is caused by reduced generalization, control animals additionally performed a click detection paradigm afterwards. D: Receiver operating characteristic: sound vs. excitation. Open symbols depict control animals; filled symbols depict animals from the experimental group. Distribution of all sessions according to the false alarm rate (y axis) and hit rate (x axis) for sound frequency discrimination (left panel) and detection of optogenetic excitation (right panel). Dashed lines represent levels of significance according to the Chi Square statistic. E: Progress in performance over all sessions. Open symbols depict control animals; filled symbols depict animals from the experimental group. Performance for detection of excitation is shown as sensitivity of the first vs. the last session for all animals individually.

## Discussion

In this study, our implant differentially stimulated two points along the ICc tonotopy and permitted using optogenetic excitation to generate behavioral responses to complex artificial cues. We tested the ability of mice to detect two independent points of optogenetic excitation within the ICc, and to discriminate between them. First, we performed a general detection task of optogenetic excitation in the ICc to evaluate the suitability of our paradigm. Seven of eight animals detected optogenetic excitation in the ICc, demonstrating the suitability of this approach (fig. 1E). Second, we investigated the behavioral sensitivity (d’) towards optogenetic excitation resulting from irradiance at two points within the ICc tonotopy (fig. 2). As shown for optogenetic excitation of rodent spiral ganglion neurons^15^ and the ICc^14^, behavioral sensitivity in our animals increased with increasing irradiance, but at both stimulation points separately. Thus optogenetic excitation can differentially stimulate the subcortical central auditory pathway. Third, we demonstrated discrimination between points in a continuous excitation task (fig. 3). To our knowledge, this study is the first systematic, long-term behavioral analysis of optogenetic excitation in the auditory pathway and the first proof-of-principle for differential and behaviorally relevant optogenetic excitation of the sub-cortical auditory pathway in freely moving, behaving animals.

### Variability of behavior as a consequence of failed generalization

In total, eight animals were injected with ChR2-AAV and implanted with two independent outlets along the tonotopy of the ICc. One animal did not generalize from sound stimulation to optogenetic excitation (overall sensitivity d’ = −0.1375, three sessions), and was additionally tested in the click-detection paradigm, a stimulus that resembles the temporal pattern of the light stimulus (data not shown) and again failed to respond. Therefore, as the implant was placed correctly, the failure in the optogenetic task was likely based on poor generalization between tasks (discrimination to detection). The remaining two animals, which successfully performed the optogenetic detection task but not the task involving differential stimulation, either had lost the implant during experimentation (M3), or developed a health problem and experiments were terminated (M2).

Despite our success in establishing differential optogenetic excitation of the ICc, we observed considerable variability between individual mice. For two animals, baseline light levels for the discrimination task had to be lowered. Since all mice were able to discriminate between tone pips of different frequencies, the limiting factor might not have been the discriminability of the optogenetic stimuli per se, but the generalizability from the acoustic to the optogenetic percept.

### Perception of optogenetic excitation versus acoustical stimulation

Although all animals that reached the final experimental stage discriminated between the two points of optogenetic excitation, our study does not elucidate the exact percept. Two factors of excitation could have contributed to the discriminability: ‘level’ and ‘(sound) frequency’ cues. Level cues could depend on a change in the overall excitation, while frequency cues would be based on a shift of excitation along the tonotopy. Perceptually, such level cues could have created a difference in the overall “loudness” of baseline and target, independent of spectral aspects. In that case, one needs to assume that stimulation at the two points resulted in widespread excitation in the entire ICc and a broad spectral percept. Alternatively, widening or narrowing the excitation area could have altered the spectral bandwidth of the percept. It is less probable that exclusively pitch differences derived from the anatomical position of the excitation points within the IC caused the discriminability of the target and baseline. In that case, discriminability between two auditory percepts differing only spectrally would have been possible without the adjustment of the baseline irradiance in animals M4 and M5.

We assume that both spectral and loudness cues contributed to the discriminability. Although the baseline amplitude had to be reduced for two animals to observe a reliable sensitivity, two other animals even performed the task with a roving of baseline amplitude and thereby a roving of irradiance for each single light pulse, disproving discrimination based on alterations in the overall ‘loudness’. Thus, different percepts elicited in the different animals, accompanied by the animals’ ability to generalize, mall ay explain the observed variability. Nevertheless, previously described scenarios were more complex, artificial stimuli that strongly differ from simple excitation at one position.

Independently of task complexity, however, the percept elicited by optogenetic stimulation may have been considerably different from that during acoustic stimulation, as reflected in reaction times. Under both conditions (discrimination and detection), optogenetic excitation was detected faster (∼70 ms) than sound stimulation. Even when considering a delay of sound transmission from the loudspeaker (∼3 ms) and the delay between ear and midbrain (∼4-5 ms ^21^), more than 60 ms remains between stimulus conditions. How can this be explained? Perhaps by a dependence of the discrimination of pure tones on auditory cortical processes, whereas optogenetic excitation triggers a cortex-independent pathway underlying the behavioral responses. Indeed, it has been previously demonstrated that alternative cortex-independent pathways exist and that they almost immediately mediate frequency discrimination after cortical lesions in mice^22^. Since the IC is crucial for the discrimination of pure tones, as its inactivation inhibited discriminability of auditory stimuli^18^, those pathways likely originate in the auditory midbrain.

Further physiological experiments revealing the responses of ICc neurons to optogenetic excitation and sound will allow assumptions concerning the artificially generated percept. Additionally, behavioral and physiological approaches should reveal whether different pathways contribute to the perception of sounds and of optogenetic stimuli.

### Suitability of the paradigm for the long-term evaluation of optogenetic excitation

Previous studies of optogenetic excitation of the auditory pathway in freely moving mice used the shuttle box paradigm^14,15^. Although the avoidance-learning procedure provides significant advantages, these paradigms cannot be easily applied to all research questions. In experiments using complex schedules requiring repetitive testing of animals, reward association positively influences behavioral outcomes^23^, and the reduction of stress and avoidance cues improves behavioral performance in mice^24^. In our paradigm, animals were not restrained during the process of attachment. reducing forces on implants and avoiding negative associations with the setup. Repetitive handling, however, made the paradigm time consuming. Nevertheless, stable behavioral performance to sound stimulation and optogenetic excitation was observed over three to six months. The different evaluations covered by our study demonstrate the flexibility and advantage of the paradigm for the long-term evaluation of optogenetic excitation in freely moving animals.

### Future optimization of optogenetic excitation of the auditory midbrain

The inferior colliculus acts as the midbrain hub for the integration of monaural and binaural ascending pathways, ascending projections to higher auditory and non-auditory areas, and, additionally, exhibits an intercollicular pathway^25–27^. Within its circuitry, approximately 25% of all neurons are GABAergic; the remaining are glutamatergic^28–31^. Thus, targeting ChR2 expressed in the IC under the control of the pancellular promotor CAG will excite both excitatory and inhibitory neurons. With regard to the potential use of optogenetics in auditory prostheses, cell-type selectivity could stimulate excitatory neuronal populations in different iso-frequency laminae without altering the entire network dynamics, thus evading a problem of electrical stimulation of the IC^32^. Although our approach did not target excitatory neurons, generalization between sound- and optogenetic excitation was rapid, and sensitivity values did not differ greatly. That the excitation of mixed neuronal populations did not negatively influence the behavioral outcome in our approach might be due to the relatively large distance between excitation points along the tonotopy (∼700 µm). Greater excitatory specificity might be required for higher spatial resolution when using stimulation at smaller distances.

What would be the limit of spatial resolution with our current approach? To estimate how the stimulation activates different areas of the IC, we developed a computational model of light activation (fig. 6A). Based on estimates of light spread^33^ and neuronal activation threshold^11^, we calculated the percentages of neurons activated at different distances from the fiber tips (fig. 6B). Up to 28.4% of optogenetically excited neurons were activated by both outlets at 6.5mW, for which we observed successful discrimination (fig 6C). Accordingly, mice can tolerate as least this level of overlap, and we accepted 29% as a conservative estimate for the upper bound for discrimination. Our detection experiments revealed that mice detected much lower light levels (fig. 2), and co-activation strongly depended on light level. Thus, we expect that light outlets can be moved considerably closer while preserving discriminability (fig. 6D). At 0.6 mW, for which 4 out of 8 stimulations resulted in behavioral detection sensitivities d’>1, stimulation at outlets spaced as close as 50 µm should be possible. But even at 1 mW, which was clearly above the detection threshold for 7 out of 8 tested outlets, a distance of 250 µm should be resolvable. If implanted in a human inferior colliculus, the distance we used in mice (700 µm) would correspond to a resolution along the tonotopic axis of ∼0.35 octaves^34^. Thus 100 µm would correspond to ∼0.05 octaves in the human IC, even without expression limited to excitatory cells. However, further experiments are necessary to test discriminability and excitatory specificity.

**Figure 6:**
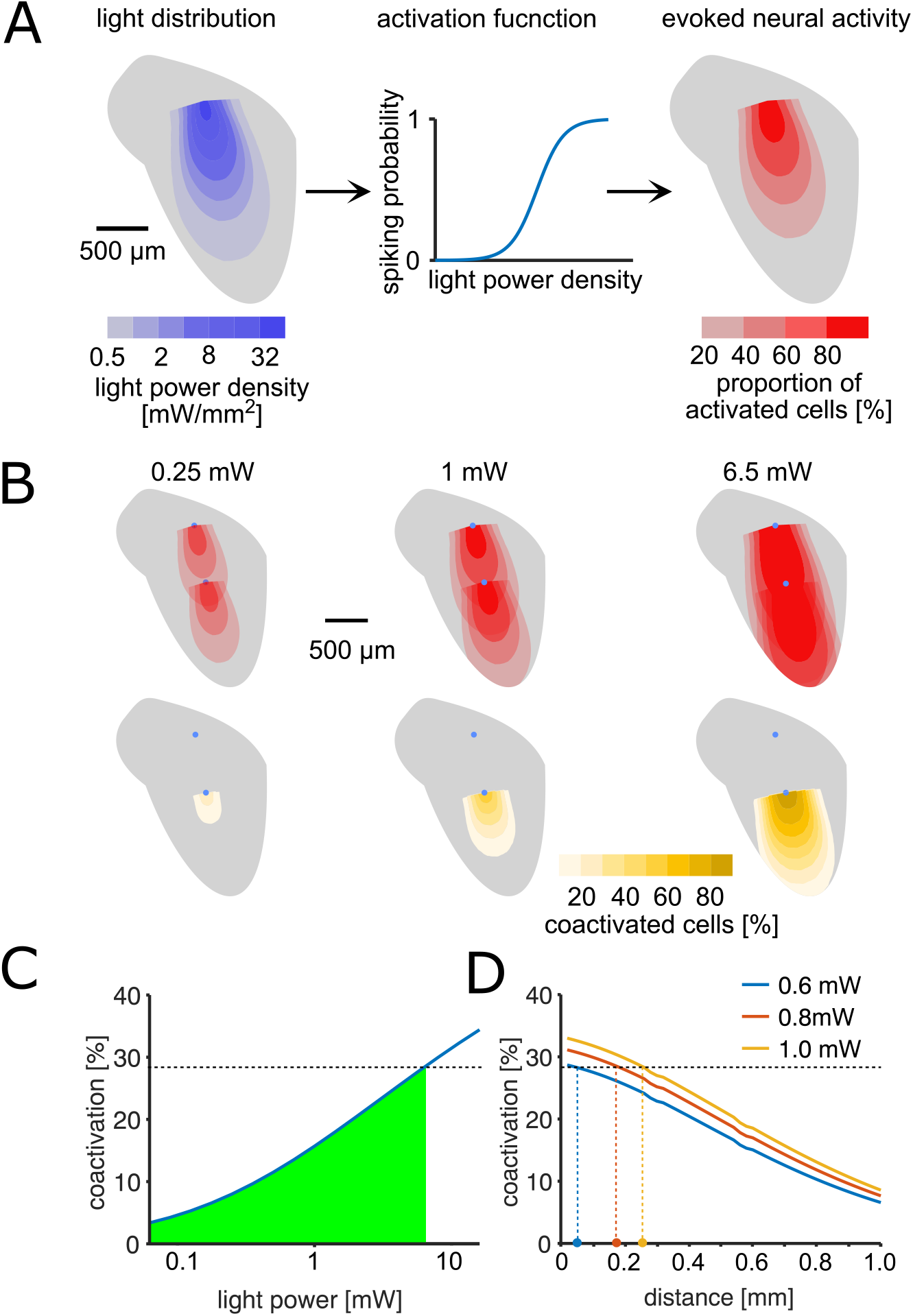
Estimation of light distribution and spread of neural activity throughout the IC. A: Model layout: light distribution was modelled using an angle-depended exponential decay of light power density^33^,^38^. Left: Color coded 2D profile of the light distribution cone estimated for a 1 mW stimulus. The estimated light power density was used to estimate the probability to fire at least one spike for neurons in each voxel (middle). The activation function was fit using data from Mattis et al. (2011). Right: 2D profile of activation cone for a l mW stimulus. B, upper row: neural activation profiles at three different output light levels for consecutive stimulation at two different outlets within the IC. The position of the fiber tips are marked with blue dots, the distance of 700µm corresponds to the setup used in our experiments. Color scale is the same as in (A). Lower row: profiles of percentage of neurons activated by both stimuli (‘coactivated’) in each voxel. C: Percentage of neurons that are activated by both outlets compared to the overall activation by a single outlet as a function of light power. The green area marks coactivation that is smaller than at 6.5 mW, for which animals were able to discriminate between stimulation at the two outputs (fig. 4). D: Estimation of minimal distance of the light outlets for lower light levels, based on the assumption that % coactivation limits the discrimination and that mice successfully discriminated the 6.5 mW stimulus with 28.5% coactivation. Colored dots at the x-axis are estimated of discriminable outlet distances at light levels at and above the detection thresholds.

Another issue is the relatively slow kinetics of ChR2. ChR2 recovery requires 10-20 milliseconds (dependent on stimulation rate and physiological conditions^35,36^). Regarding the potential of optogenetics for restoring auditory function, ChR2 might not be a suitable ion channel for precisely timed processing; other channels such as Chronos should achieve more precise coding of auditory information^14,37^. However, while the use of Chronos produced more synchronized neuronal responses in the mouse IC at rapid stimulation rates, it did not improve behavioral detectability compared to ChR2^14^. It remains to be seen whether ultra-fast ion channels can improve sensitivity for more complex optogenetic excitation cues.

## Perspective

The present study demonstrates that differential stimulation of the central auditory system is possible using optogenetics and that the auditory midbrain represents a promising target for the generation of complex, continuous sensory patterns. Further research is needed to reveal the actual network activity resulting from differential optogenetic excitation of the sub-cortical auditory pathway. Behaviorally, it remains to be shown how the discriminability of two neuronal ensembles depends on the distance of excitation outlets and the limits of spatial resolutions.

Our paradigm could be very helpful for future work on optogenetic restoration of sensory function, since it provides a high level of flexibility and a reliable behavioral read-out of optogenetic excitation over periods of several months. Our results demonstrate that optogenetic excitation of sub-cortical sensory nuclei could be a powerful tool to restore sensory function in the auditory system and beyond.

## Material & Methods

### Animals

All experiments were performed in accordance with the animal welfare regulations of Lower Saxony and with the approval from the local authorities (State Office for Consumer Protection and Food Safety / LAVES, permission number 33.9-43502-04-18/2802).

In total, 11 adult male mice, bred at the University of Oldenburg animal facilities, were used. All mice had a C57BL/6.CAST-Cdh23^*Ahl+*^ background (the Jackson Laboratory, #002756). Standardized cages were used for housing, equipped with cage enrichment. Animals were single housed but with visual and olfactory contact to neighboring animals at a reversed 12/12 hour dark-light cycle. Experiments were conducted during the dark period for 1 to 3 times/day. During experimental periods, animals had unlimited access to water but were food-deprived to a moderate extent (85 - 90 % of their ad libitum weight). Weight and wellbeing was scored daily.

### Implants and surgery

#### Implants

The optogenetic auditory midbrain implant (oAMI) consisted of two separate fiber outlets with a difference in depth of 700 µm, terminating at two different positions with the right IC (dorsal and ventral, see Fig. 1B-D). PlexBright fiber stub implants were used with LC ceramic ferrules (1.25 mm outer diameter, 6.45 mm length), equipped with 1.2 cm long high performance optical fibers of a diameter of 110/125 µm and a numerical aperture (NA) of 0.66 (Plexon Inc, USA). At the distal region of the ferrule, a small notch was cut with a diamond plate grinder for a better grip of the dental cement (Vertex Self-Curing, Vertex Dental, Netherlands). Prior to attachment, fibers and ferrules were cleaned with dust free wipes and isopropanol. Fiber stubs were aligned with a distance of 700 µm under a microscope and covered with dental cement, from the medial part of the fibers leaving ∼1/3 exposed, to the medial part of the ferrules. Both outlets were tested for their functionality and stored in dust-free plastic boxes until implantation.

Implant position and virus expression were evaluated in frozen sections (40 µm) of perfused and coronal cut brains. Slices were additionally stained with DAPI (1 µg/ml, D9542, Lot #096M4014V, Sigma, USA).

#### Transfection & implantation

Mice had unlimited access to food for at least three days prior to surgery. Anesthesia was induced using 2% isoflurane (CP Pharma, Germany) provided with oxygen and a flowrate of approx. 0.9 l/min. Animals received 0.1 mg/kg meloxicam (Metacam, 2 mg/ml, Boehringer Ingelheim, Germany) in Ringer-lactate solution subcutaneously. Surgery was performed on a heating pad in a stereotactic frame for rodents (Model 900, Kopf Instruments, USA) using zygomatic bars. The head was fixated with an angle of 7° in the rostrocaudal axis to reach the same z-position for Bregma and IC (see Fig. 1B). After proper fixation, anesthesia was reduced to the maintenance dose of 1.2-1.5 %. A small incision was made from the caudal end of the frontal bone near the coronal suture and extended caudally until full exposure of the interparietal bone. The periost was removed and 4-5 holes were drilled into the parietal bones and equipped with autoclaved screws. Bregma and Lambda were determined and a small trepanation was drilled above the right IC (X= 0.5; Y= −5.2, relative to Bregma). The viral construct (rAAV5-CAG-ChR2-GFP, 2.9*10^12^ molecules/ml, Lot# AV4597D, UNC GTC Vector Core, USA) was injected using a 10 µl syringe and an ultra-micro pump (UMP3 with SYS-Micro4 Controller, WPI, UK) and 35 G needles on a micron-resolution manipulator. 3 x 300 nl of the construct were injected at an angle of −26° relative to Bregma (Z^1^= −1.5, Z^2^= −2.1, Z^3^= −2.7) with a rate of 150 nl/min. After each injection, the needle was maintained at its position for five minutes. Implantation was performed using a custom made tool adjusted on a standard cannula holder on the manipulator using the same X and Y coordinates as for the injection, with the ventral outlet as reference. The oAMI was implanted at Z= −2.4 (Fig. 1B-D), the craniotomy was covered with sterile Vaseline and the implant and screws were completely covered with dental cement. A second dose of meloxicam in Ringer solution was administered subcutaneously. Throughout the surgery and a postsurgical period of 2-3 hours, breathing frequency and wellbeing were monitored and the temperature was kept at ∼37°C.

During a recovery period of two weeks, animals received 1.77 mg/kg meloxicam (Metacam, 0.5 mg/ml, Boehringer Ingelheim, Germany) orally, mixed with a high-energy nutritional supplement (DietGel Boost, ClearH2O, USA) every 12 hours for at least seven days. During the whole period, animals had unlimited access to water and food.

### Behavioral experiments

#### Behavioral setup & paradigm

All behavioral experiments were performed using the following reward-based operant Go/No-Go paradigm.

Animals were placed on an elevated circular shaped wire mesh runway, lined with outer and inner walls of Plexiglas (outer Ø = 29 cm, inner Ø = 20 cm, height = 27 cm, fig 5.1A) positioned in a double-walled sound attenuated booth with pyramid foam covered walls (Industrial Acoustics Company GmbH, Germany). One side of the runway was equipped with a small platform (5 x 3 x 4 cm). Once the animals ascended the platform, which was detected by a light barrier, a random waiting time started, ranging from 1.25 to 5.25 s in steps of 1 s, followed by a target presentation. The onset of the target triggered a 1 s response window. If the animals descended from the platform within the window (,go’), a food pellet (0.02 g, Dustless precision pellets rodent, grain based, #F0163, Bio-Serv, USA) was delivered at the opposite side of the runway by a feeder (hit). If the animals did not leave the platform (miss), a new trial was presented after a newly drawn waiting time. In order to estimate the amount of coincidently correct responded trials, 27.27 % were sham trials (trials without stimulus presentation), pseudo-randomly integrated into each session. These trials had the same distribution of waiting times as the target trials and neither a response to (false alarm), nor a correct rejection of the sham trials were punished or rewarded. A session contained 40 trials and 15 sham trials and usually lasted 15-30 minutes.

To reduce the possibility of visual detection of the light stimulation, the booth was additionally equipped with a blue light LED strip at the ceiling (57 cm, 30 LEDs). This masking LED was continuously pulsing in the same rate as the chosen stimulation task during every single session, also during sound training and experiments.

Behavioral experiments were controlled by a custom Software (Github repository: https://github.com/Spunc/PsychDetect), written in MATLAB (The Mathworks, RRID:SCR_001622). Pellet dispenser and light barriers were custom made (University of Oldenburg workshop) and controlled by a microcontroller (Arduino UNO, Arduino AG, Italy) connected to a Windows PC. Experiments were carried out in darkness and were observed visually by a camera with infrared LEDs.

#### Sound stimulation

For sound presentation, a speaker (Vifa XT 300/K4, Denmark) was placed at the ceiling of the booth, approx. 0.8 m above the platform. Sound was generated using a high-fidelity sound card (Fireface UC, RME, Germany) connected to the PC. Sound was played back at 96 kHz sampling rate. The speaker was calibrated on the platform at the approximate position of the head of the animals using a measurement microphone (model 40BF, G.R.A.S, Denmark).

### Sound frequency discrimination task

Animals had to report a change in frequency within a continuous sequence of tone pips, presented with a rate of 5 Hz with a roving of tone levels between 60 and 66 dB SPL (randomly drawn) to avoid the detection of differences in loudness when the shift in frequency occurred. The shift was +0.241 octaves relative to the baseline frequency, which was between 10-18 kHz in steps of 2 kHz, presenting one frequency per session, randomly chosen (fig 5.3A).

### Handling & auditory training

Upon arrival from the local animal facility at the age of 9 to 12 weeks, each animal was kept for at least two days without handling and food deprivation to habituate to the novel situation of single-housing, cage enrichment and inverted dark-light cycle. After food restriction, mice underwent a strict handling protocol prior to experiments, performed for one week two times per day for at least 15 minutes to reduce avoidance behavior. Animals were habituated to the experimental setup in silence and darkness two times per day for four times (2 x 10, 1 x 15, 1 x 20 min), provided with pellets placed in the feeder bowl. Auditory training in the sound frequency discrimination task at a baseline frequency of 10 kHz was conducted two times per day with a minimum of 1.5 h in between, introduced with relatively short waiting times (0.2-0.7 s) and without sham trials. The waiting time was gradually increased until a stable performance at waiting times between 1.25 and 5.25 s, which usually lasted 3-7 days, was observed in several consecutive sessions. A training session was terminated after 30 minutes or 40 received rewards.

#### Optogenetic excitation

Two 465 nm compact LED Modules on a dual LED commutator (PlexBright, Plexon Inc, USA) were placed at the ceiling of the booth, approx. 1 m above the center of the circular runway. LEDs were independently connected to LED drivers (LEDD1B T-Cube, Thorlabs, USA), on which the maximum current was set to 200 mA. As sound cues, light pulses were generated using the sound card connected to the PC.

Light was delivered using optical patch cables (0.66 NA; PlexBright, high-performance fibers and LC ceramic ferrules, Plexon Inc, USA). Ceramic sleeves were used to connect cables and implants. The functionality of each cable was tested prior to each session. Stubs were gently connected without fixation of the implant or the head of the animal and covered with a black tube to additionally avoid visual detection of light stimulation.

Level of light output at corresponding voltage values of our system were verified at the fiber stub tips using a digital optical power and energy meter (PM100D, Thorlabs, USA) in combination with a photodiode power sensor (S121C, Thorlabs, USA) and an oscilloscope.

### Simple detection of optogenetic excitation

To evaluate the usability of our paradigm and to test for the ability of 2-point detection, animals performed a simple detection task of a short pulse train in silence for one outlet (number of pulses: 4; pulse duration: 0.1 ms; rate: 13.8 Hz; light level: ∼6.5 mW). After achieving a high performance (d’ > 1.2) for at least two sessions, the level for light was varied (∼0.6; 1; 2; 6.5 mW). If a significant detectability was still observable for the lowest light level, another combination of light levels was presented (∼0.09; 0.2; 1.5; 3.9; 5.2 mW) until reaching a subthreshold level for that stimulation point (non-significant according to the Chi Squared test). Each light level was presented at least 20 times (two independent sessions). These experiments were then repeated for the respective other outlet. The whole detection period usually lasted 2-3 weeks.

### 2-point discrimination task

The animals had to report a change of stimulation point within a continuous sequence of light pulses (number of pulses: 2; pulse duration: 0.1 ms; rate: 5 Hz). Usually, the ventral outlet represented the baseline point of excitation and each pulse (target and baseline) was presented with a level of ∼2.2 mW, if not stated otherwise.

If a stable performance could not be observed after three consecutive sessions for an equal light level in both points, the level of the baseline stimulation point was reduced (25, 50 %). The period for these experiments strongly depended on the performance of each individual, lasting from 1 week (M4) to 4 months (M6).

### Control task

To investigate if a visual cue leads to a significant performance in our optogenetic experiments, a control task was implemented, partly adapted from Wrobel and colleagues (2018). Animals had to perform the simple detection task in five consecutive sessions with the highest light level (∼6.5 mW); targets appeared with 50 % probability in either one of the two outlets. In the second session, the light path of the dorsal outlet was blocked using black sponge rubber at the connector of stub and cable. During the third session, the other path was blocked with the same procedure whereas the other sessions were conducted without blockage. During the blocked sessions, the light stimulus in the blocked point should trigger the same response as in the unblocked if the target is perceived as a visual cue from reflections at the implant or the chamber. This period lasted 2.5 days.

### Control animals

To control for the necessity of the opsin, 2 animals have been trained, transfected and implanted as the other remaining animals, but were injected with a control construct, encoding only GFP (rAAV5-CAG-GFP). Experiments were conducted in the same order as for most of the experimental animals (M1-M5): sound frequency discrimination (including sessions were animals were attached to patch cables for habituation); followed by detection of optogenetic excitation. In the latter, control animals were repetitively tested in six consecutive sessions. For the control animals, a click detection task was conducted within the next step to test whether these animals were able to generalize between cues.

### Data analysis and statistics

In all experiments, for each session *i* and stimulus class *s*, the sensitivity d’ was calculated as:

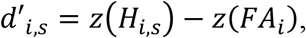

where *z()* is the inverse of normal cumulative function, *H*_*i,s*_ is the hit rate for the stimuli with parameters *s* in the *i*th session *P(response*|*stimulus s)* and *FA*_*i*_ is the false alarm rate *P(response*|*sham)*.

To discriminate between significant and non-significant detection, a Chi-Square (*x*^2^) statistic for the 2 x 2 contingency table (‘go’ and ‘no-go’ responses during trials and sham trials) was calculated as:

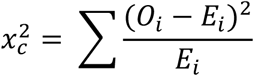

where c is the number of degrees of freedom, *O_i_* are the observed and *E_i_* the expected values for the *i*th session.

### Model for spread of neural activation

In order to estimate the overlap of neural activation for two stimulation sites, we conceived a 3D model for light spread and neural activation at each voxel. The light spread was modelled as an exponential decay of the light power density *p* along the fiber axis *z* ^38^:

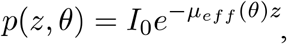

where *I*_*0*_ is the respective power density at the fiber tip and *µ*_*eff*_*(*θ*)* the effective decay constant at the angle *θ*. *µ*_*eff*_ was estimated separately for each angle *θ* using data from Gysbrechts et al. (2016). The resolution of the model was 10×10×10µm voxel, with corresponding *θ* and *µ*_*eff*_l. We defined neural activation of a voxel as the proportion of neurons within the voxel that fired as least one spike in response to the light stimulus. We used data for the light-dependence of photo currents and spike probability for ChR2 from Mattis et al. (2011) to estimate the activation *a* at each voxel, depending on the light power density *p*:

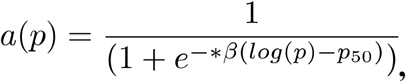

with the slope parameter *β = 0*.*2624* and 50% activation at the power density *p*_*50*_ *= 0*.*814 mW/mm*^*2*^. The activation *a* can be interpreted both as the probability of a neuron in a given voxel to fire at least one spike or as the number of neurons in the voxel to responding to the stimulus.

Using the voxel-resolved neural activation, we could estimate the number of ‘coactivated’ neurons, defined as the number of neurons that would be active after stimulation from either of the two simulated outputs as the joined probability (or activation) of a neuron to fire at least one spike in response to both outlet 1 and 2: *a*_*1*_*\*a*_*2*_. By varying output light power and distance between light outlets we could simulate the relative amount of coactivated neurons by taking the sum over all voxels:

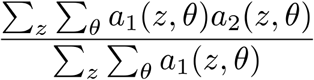

## Acknowledgements

The authors thank P.F. Apostolides, G. A. Manley, and G. Quass for the helpful comments and the productive scientific discussions. Additionally, the authors would like to thank J. M. Sleeboom, K. Schulze, A. A. Korte, and Y. Karsten for the support with animal handling and behavioral training, L. Torres Mapa for acquiring the detailed implant image in figure 1 and www.stels-ol.de for proofreading and language correction of the manuscript.

This work was supported by the DFG Cluster of Excellence ‘Hearing4all’ (grant no. EXC 1077/1 & EXC 2177/1).

## References

1. NIDCD. Cochlear Implants. NIDCD https://www.nidcd.nih.gov/health/cochlear-implants (2015).

2. Fan-Gang, Z., Rebscher, S., Harrison, W., Xiaoan Sun, & Haihong Feng. Cochlear Implants: System Design, Integration, and Evaluation. IEEE Rev. Biomed. Eng. 1, 115–142 (2008).

3. Middlebrooks, J. C., Bierer, J. A. & Snyder, R. L. Cochlear implants: the view from the brain. Curr. Opin. Neurobiol. 15, 488–493 (2005).

4. Colletti, V., Shannon, R., Carner, M., Veronese, S. & Colletti, L. Outcomes in Nontumor Adults Fitted With the Auditory Brainstem Implant: 10 Years’ Experience. Otol. Neurotol. 30, 614–618 (2009).

5. Lim, H. H. & Lenarz, T. Auditory midbrain implant: Research and development towards a second clinical trial. Hear. Res. 322, 212–223 (2015).

6. Hernandez, V. H. et al. Optogenetic stimulation of the auditory pathway. J. Clin. Invest. 124, 1114–1129 (2014).

7. Kral, A., Hartmann, R., Mortazavi, D. & Klinke, R. Spatial resolution of cochlear implants: the electrical ¢eld and excitation of auditory a¡erents. Hear. Res. 18 (1998).

8. Bernstein, J. G. & Boyden, E. S. Optogenetic tools for analyzing the neural circuits of behavior. Trends Cogn. Sci. 15, 592–600 (2011).

9. Delbeke, J., Hoffman, L., Mols, K., Braeken, D. & Prodanov, D. And Then There Was Light: Perspectives of Optogenetics for Deep Brain Stimulation and Neuromodulation. Front. Neurosci. 11, 663 (2017).

10. Fenno, L., Yizhar, O. & Deisseroth, K. The Development and Application of Optogenetics. Annu. Rev. Neurosci. 34, 389–412 (2011).

11. Mattis, J. et al. Principles for applying optogenetic tools derived from direct comparative analysis of microbial opsins. Nat. Methods 9, 159–172 (2012).

12. Moser, T. Optogenetic stimulation of the auditory pathway for research and future prosthetics. Curr. Opin. Neurobiol. 34, 29–36 (2015).

13. Moser, T. & Dieter, A. Towards optogenetic approaches for hearing restoration. Biochem. Biophys. Res. Commun. (2020) doi:10.1016/j.bbrc.2019.12.126.

14. Guo, W. et al. Hearing the light: neural and perceptual encoding of optogenetic stimulation in the central auditory pathway. Sci. Rep. 5, 10319 (2015).

15. Wrobel, C. et al. Optogenetic stimulation of cochlear neurons activates the auditory pathway and restores auditory-driven behavior in deaf adult gerbils. Sci. Transl. Med. 10, eaao0540 (2018).

16. Gill, J. V. et al. Precise Holographic Manipulation of Olfactory Circuits Reveals Coding Features Determining Perceptual Detection. Neuron 108, 382-393.e5 (2020).

17. May, T. et al. Detection of Optogenetic Stimulation in Somatosensory Cortex by Non-Human Primates - Towards Artificial Tactile Sensation. PLoS ONE 9, e114529 (2014).

18. Ceballo, S., Piwkowska, Z., Bourg, J., Daret, A. & Bathellier, B. Targeted Cortical Manipulation of Auditory Perception. Neuron 104, 1168-1179.e5 (2019).

19. Marshel, J. H. et al. Cortical layer–specific critical dynamics triggering perception. Science 365, eaaw5202 (2019).

20. Owen, S. F., Liu, M. H. & Kreitzer, A. C. Thermal constraints on in vivo optogenetic manipulations. Nat. Neurosci. 22, 1061–1065 (2019).

21. Land, R., Burghard, A. & Kral, A. The contribution of inferior colliculus activity to the auditory brainstem response (ABR) in mice. Hear. Res. 341, 109–118 (2016).

22. O’Sullivan, C., Weible, A. P. & Wehr, M. Auditory Cortex Contributes to Discrimination of Pure Tones. eneuro 6, ENEURO.0340-19.2019 (2019).

23. Patterson-Kane, E., Pittman, M. & Pajor, E. Operant animal welfare: productive approaches and persistent difficulties. 11 (2008).

24. Havenith, M. N. et al. The Virtual-Environment-Foraging Task enables rapid training and single-trial metrics of rule acquisition and reversal in head-fixed mice. Sci. Rep. 9, 4790 (2019).

25. Gruters, K. G. & Groh, J. M. Sounds and beyond: multisensory and other non-auditory signals in the inferior colliculus. Front. Neural Circuits 6, (2012).

26. Malmierca, M. S. The Inferior Colliculus: A Center for Convergence of Ascending and Descending Auditory Information. Neuroembryology Aging 3, 215–229 (2004).

27. Syka, J., Popelář, J., Kvašňák, E. & Astl, J. Response properties of neurons in the central nucleus and external and dorsal cortices of the inferior colliculus in guinea pig. Exp. Brain Res. 133, 254–266 (2000).

28. Beebe, N. L., Young, J. W., Mellott, J. G. & Schofield, B. R. Extracellular Molecular Markers and Soma Size of Inhibitory Neurons: Evidence for Four Subtypes of GABAergic Cells in the Inferior Colliculus. J. Neurosci. 36, 3988–3999 (2016).

29. Ito, T., Bishop, D. C. & Oliver, D. L. Expression of glutamate and inhibitory amino acid vesicular transporters in the rodent auditory brainstem. J. Comp. Neurol. 519, 316–340 (2011).

30. Merchán, M., Aguilar, L. A., Lopez-Poveda, E. A. & Malmierca, M. S. The inferior colliculus of the rat: Quantitative immunocytochemical study of GABA and glycine. Neuroscience 136, 907–925 (2005).

31. Naumov, V., Heyd, J., de Arnal, F. & Koch, U. Analysis of excitatory and inhibitory neuron types in the inferior colliculus based on I h properties. J. Neurophysiol. 121, 2126–2139 (2019).

32. Quass, G. L., Kurt, S., Hildebrandt, K. J. & Kral, A. Electrical stimulation of the midbrain excites the auditory cortex asymmetrically. Brain Stimulat. 11, 1161–1174 (2018).

33. Gysbrechts, B. et al. Light distribution and thermal effects in the rat brain under optogenetic stimulation. J. Biophotonics 9, 576–585 (2016).

34. Ress, D. & Chandrasekaran, B. Tonotopic Organization in the Depth of Human Inferior Colliculus. Front. Hum. Neurosci. 7, (2013).

35. Grossman, N. et al. The spatial pattern of light determines the kinetics and modulates backpropagation of optogenetic action potentials. J. Comput. Neurosci. 34, 477–488 (2013).

36. Schneider, F., Grimm, C. & Hegemann, P. Biophysics of Channelrhodopsin. Annu. Rev. Biophys. 44, 167–186 (2015).

37. Keppeler, D. et al. Ultrafast optogenetic stimulation of the auditory pathway by targeting-optimized Chronos. EMBO J. 37, (2018).

38. Al-Juboori, S. I. et al. Light Scattering Properties Vary across Different Regions of the Adult Mouse Brain. PLOS ONE 8, e67626 (2013).

